# Stable modulation of neuronal oscillations produced through brain-machine equilibrium

**DOI:** 10.1101/2021.10.14.464387

**Authors:** Colin G. McNamara, Max Rothwell, Andrew Sharott

**Affiliations:** MRC Brain Network Dynamics Unit, Nuffield Department of Clinical Neurosciences, University of Oxford, Oxford OX1 3TH, United Kingdom

## Abstract

Brain stimulation is predominantly delivered independent of ongoing activity, whereas closed-loop systems can modify activity through interaction. They have the unrealised potential to continuously bind external stimulation to specific dynamics of a neural circuit. Such manipulations are particularly suited to rhythmic activities, where neuronal activity is organised in oscillatory cycles. Here, we developed a fast algorithm that responds on a cycle-by-cycle basis to stimulate basal ganglia nuclei at predetermined phases of successive cortical beta cycles in parkinsonian rats. Using this approach, we demonstrate a stable brain-machine interaction. An equilibrium emerged between the modified brain signal and feedback-dependent stimulation pattern, which led to sustained amplification or suppression of the oscillation. Sustained beta amplification slowed movement speed by altering the mode of locomotion. Integrating an external stimulus with network activity in this way could be used to correct maladaptive activities and to define the role of oscillations in fundamental brain functions.

## Introduction

Oscillatory activity in the brain is supported by reciprocal connectivity that allows the firing of action potentials in connected areas to be precisely timed in relation to each other at specific frequencies (Buzsáki and Draguhn, 2004). When recorded in the local field potential or electrocorticogram (ECoG), such activities typically wax and wane in amplitude, indicating the fluctuating strength and phase-stability of these oscillatory interactions over time (Buzsáki et al., 2012; Feingold et al., 2015). Achieving precise manipulation of neuronal oscillations is of primary importance, given their ubiquitous association with widespread brain functions (Buzsáki *et al*., 2012) and role in many prominent brain disorders including schizophrenia, depression, motor disorders, Alzheimer’s disease, addiction and epilepsy (Schnitzler and Gross, 2005; Uhlhaas and Singer, 2006). Simply disrupting a relevant network node will heavily affect many unrelated processes and does not allow the oscillation to be manipulated bidirectionally. An alternative approach is to deliver a perturbation on a specific phase of the oscillation (Siegle and Wilson, 2014), which can energise or dampen ongoing activity (Cagnan et al., 2017; Holt et al., 2019; Kanta et al., 2019; Nicholson et al., 2018; Peles et al., 2020; Rosin et al., 2011; Takeuchi et al., 2021). Hitherto, such approaches have been defined by the effect of the perturbation on the amplitude or phase of input oscillation. However, if the parameters of the closed-loop allow rapid response to these perturbations, a bidirectional interaction could develop between signal and timing of stimulation. Such a fast-acting system, continuously pushing a brain network towards a desired state, has wide-ranging potential to provide physiologically relevant manipulations. The interaction should be sufficiently light touch that the brain network is still free to exhibit relevant physiological fluctuations. Using parkinsonian beta oscillations as a model, we demonstrate that a closed-loop system delivering stimulation to the basal ganglia at specific phases of a cortical oscillation can generate such brain-machine equilibrium and produce a state-like change in ongoing behaviour.

## Results

### Real-time phase tracking and phase-locked stimulation

To enable the sustained application of phase-dependent stimulation, we developed a novel approach for continuous real-time phase estimation with zero filter delay (C.G.M. and A.S. patent application WO/2020/183152), which we implemented as a digital circuit using the existing hardware of a commercially available recording system. Ordinarily, to produce such an online phase estimate the signal would be filtered with a pass band filter before a phase estimation step such as the Hilbert transform or the detection of zero crossings (Brittain et al., 2013; Cagnan *et al*., 2017; Escobar Sanabria et al., 2020; Kanta *et al*., 2019; Peles *et al*., 2020; Schreglmann et al., 2021; Siegle and Wilson, 2014; Takeuchi *et al*., 2021; Widge et al., 2018; Zanos et al., 2018). Such filters exhibit a filter delay, whereby the estimate at a given time point pertains to a fixed time in the past. While easily corrected in offline analysis, when calculating in real-time, this results in a delay that typically can amount to half a cycle or more. This delay (which is due to the algorithm, not the hardware on which it is implemented) coupled with loss of signal around stimulation artefacts and further delays due to implementation hardware represent challenges in realising a system that can react responsively on a cycle-by-cycle basis. Our algorithm operates directly on the wide band signal and provides a phase estimate for each sample with zero filter delay (see methods). Here, implementation as a digital circuit in the low-level hardware of the recording system ensured associated delays were insignificant. Together these innovations enabled the continuous delivery of phase-targeted stimulation with each stimulation informed by the immediately preceding cycle.

To demonstrate the utility of our approach, we performed phase-locked stimulation of the pathologically-exaggerated beta oscillations (Figures 1a and S1) that occur in the parkinsonian cortico-basal ganglia network (Hammond et al., 2007). Electrical stimulation (50-70 μA) was delivered to the globus pallidus (GPe) of 6-Hydroxydopamine (6-OHDA) hemi-lesioned rats at 8 equally spaced target phases of the ongoing ECoG beta oscillation. The GPe is a key node in the generation of parkinsonian beta oscillations (Crompe et al., 2020; Mallet et al., 2008). Across all phases, 58.66 ± 4.71 % (mean ± SD, n = 13 rats) of the stimuli were delivered within a quarter of a cycle of the target phase. This distribution was almost identical to that when stimulation was disabled (p = 0.093, Kolmogorov-Smirnov test, KS = 0.22; Figure S1c), suggesting phase tracking accuracy was maintained in the presence of stimulation.

**Figure 1.**
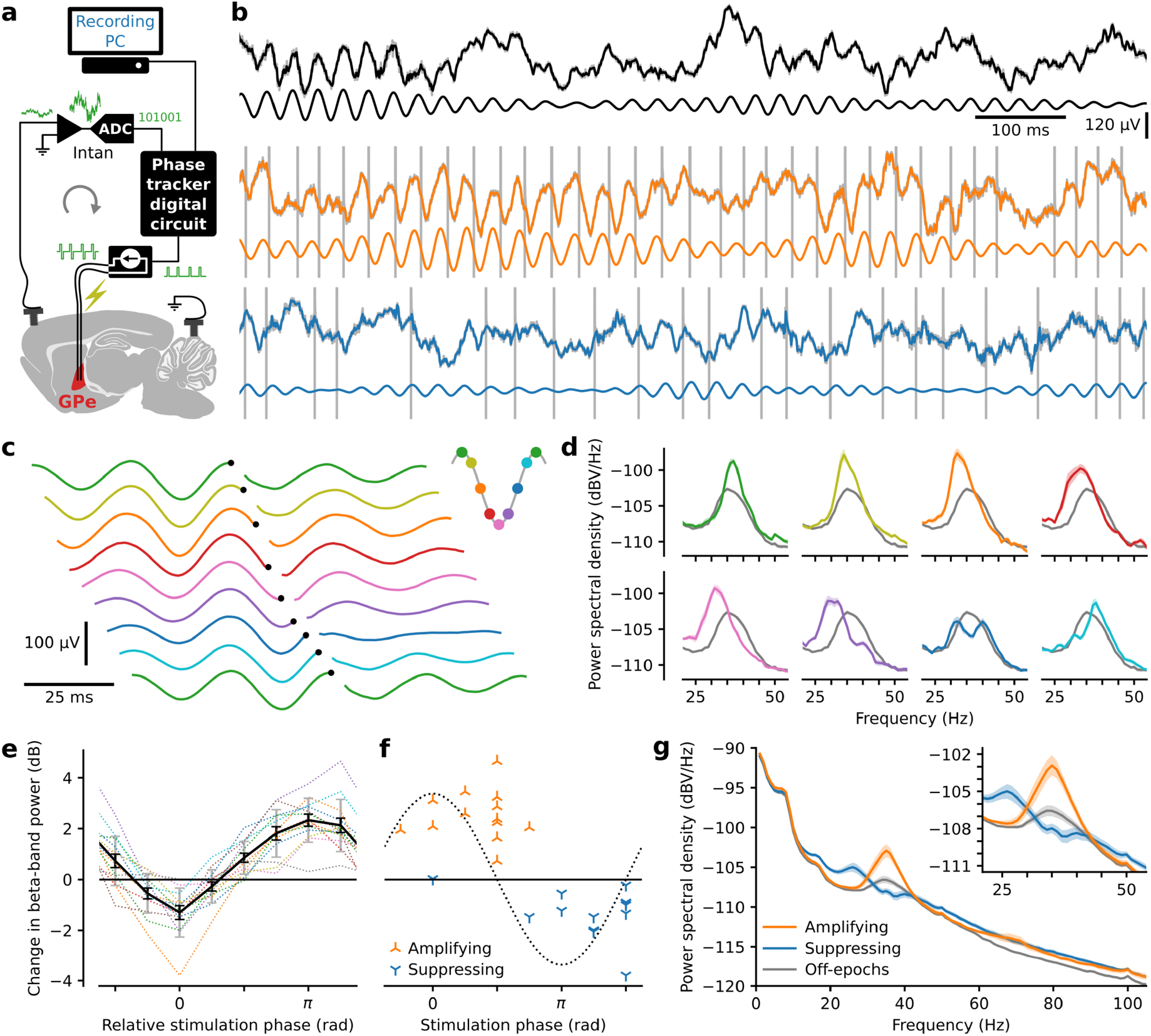
Phase-dependent modulation of beta-frequency oscillations. **a**, A digital circuit triggered delivery of electrical pulses to the globus pallidus at a predetermined beta-band ECoG phase. **b**, Example one second pre-artefact removal raw ECoG traces in awake freely behaving rats (grey) with artefact removed down sampled traces superimposed and beta-band filtered traces below. Three conditions are shown; no stimulation (top), stimulation targeted to mid-descending phase producing amplification (middle) and targeted to the mid-ascending phase producing suppression (bottom). **c**, Stimulation triggered averages from the same example block of ECoG recordings, targeted to eight equally spaced phases. To aid visualisation, trigger times (black dots) are staggered across traces and the first trace is repeated. **d**, Power spectra from stimulation on-epochs for each target phase (colours as in **c**) from the same example recordings and for all off-epochs imbedded in those same recordings (grey). **e**, Change in beta-band power (on-compared to off-epochs) due to stimulation across 13 rats for 8 target phases aligned by the most suppressing target phase for each rat. Dotted lines show the values for each rat with mean ± SEM in black and standard deviation in grey. **f**, Absolute phase versus beta-band power of the most suppressing and most amplifying target phases for each rat. The dotted line represents the beta-cycle. g, Power spectral density plots (mean ± SEM, n=13 rats).

### Phase-dependent modulation of oscillatory power

It was apparent in the raw data that stimulation at some target phases reduced beta-band activity whereas others increased it (Figures 1b-d). Absolute target phase modulated beta-band power (p = 1.8e-07, Kruskal-Wallis test, H_7_ = 44.44) and the phases of maximum amplification (0.38 ± 0.27 π rad) and suppression (1.35 ± 0.30 π rad) were different (p = 1.2e-07, Watson-Williams test, F_1, 24_ = 55.08), occurring approximately anti-phase (Figures 1e and f). Maximally amplifying (2.54 ± 0.93 dB) and suppressing (−1.30 ± 0.93 dB) stimulation enhanced and flattened the beta-band spectral peak, respectively (Figure 1g). Similar modulation could be achieved through stimulation of STN (Figure S2).

### An equilibrium emerged between the stimulus train and the oscillation which depended on stimulation phase

Modulation of beta-band oscillatory power represents a change to the signal that the algorithm uses to calculate phase. Next, we addressed the hypothesis that the feedback loop would produce an interplay between the temporal properties of the oscillation and stimulus train, which would differ between amplification and suppression. The pattern of stimulation was visibly different between these conditions (Figures 1b and 2a), as were the inter-stimulus interval histograms and autocorrelograms (Figures 2b-e). We thus investigated if there was a matching difference in the temporal structure of the fluctuations in oscillatory amplitude. Coefficient of variation-type measures when applied to time intervals describe variation in time. However, when directly applied to constant sample rate processes such as oscillatory amplitude, they describe variability in amplitude, not time. Thus, to describe the temporal structure of the oscillations, we developed the Temporal Variation Index (TVI), which is the standard deviation of the derivative of the Hilbert amplitude envelope divided by the envelope mean (see methods and Figure S3).

**Figure 2.**
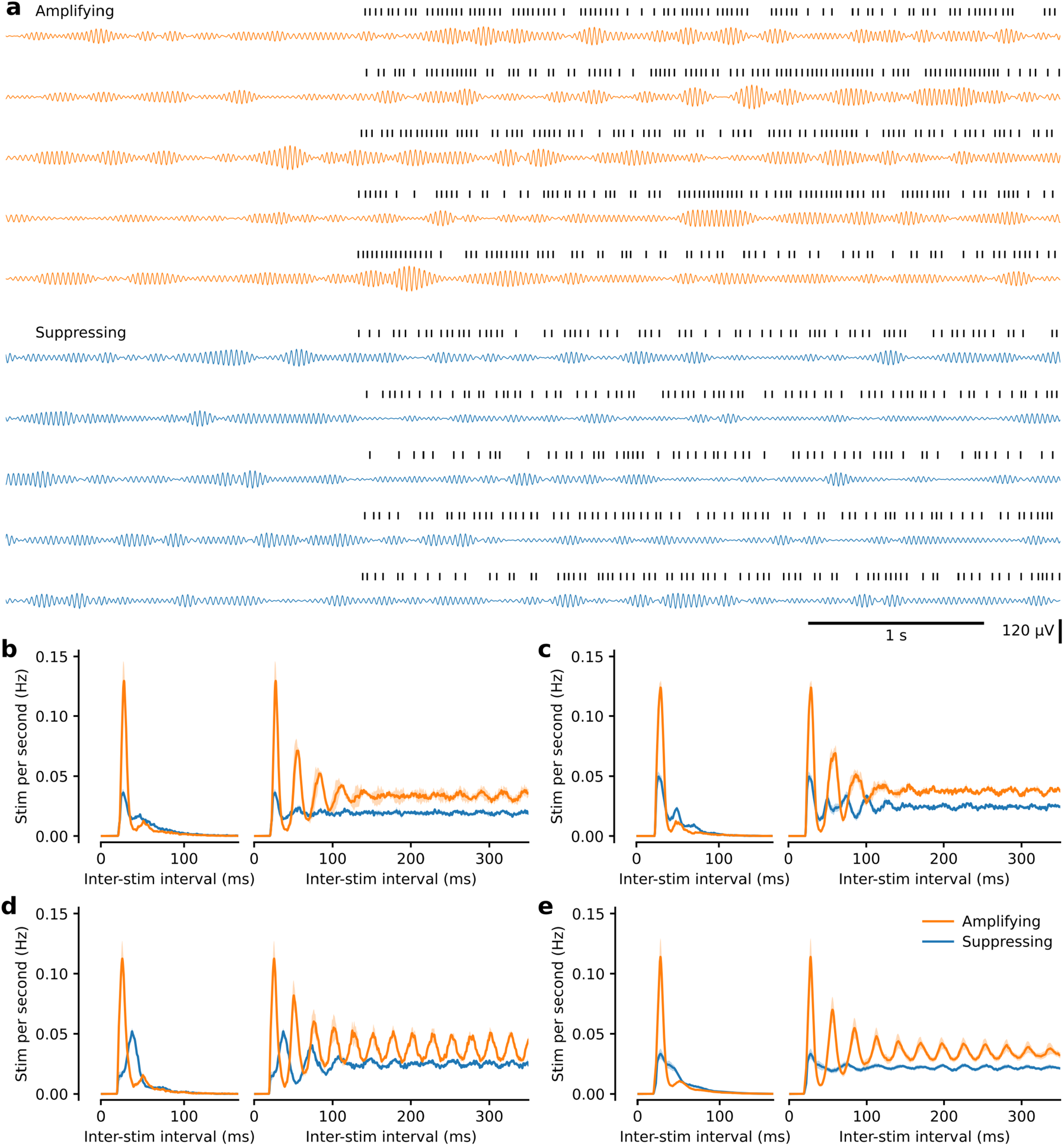
The pattern of stimulation differed between amplifying and suppressing stimulation. **a**, Example beta-filtered ECoG traces from maximally amplifying and suppressing conditions from the same block. Five consecutive off-to on-epoch transitions in each case. Stimulation times are shown above each trace. Note the difference in stimulation pattern between the two conditions. Beta power is similar during both sets of off-epochs, is higher during stimulation targeted to an amplifying phase and lower during suppressing stimulation, yet still waxes and wanes in all conditions. **b**, Distribution of inter-stimulation intervals to the next stimulation (left) and to all subsequent stimulations (right; autocorrelogram) from the same rat as **a. c, d**, Same as **b** for two additional example rats. **e**, Same as **b** for all animals (mean ± SEM, n = 13 rats). Note the only experimenter-controlled variable change between amplifying and suppressing conditions was selection of a different target phase.

The amplified oscillation had a lower TVI than the suppressed oscillation indicating it was more stable over time (p = 0.0005, Wilcoxon signed-rank test (WSRT), W_13_ = 1; Figure 3a). In line with this, the variation in short timescale dynamics of the stimulation train was lower during amplifying stimulation (CV_2_; p = 0.0005, WSRT, W_13_ = 1; Figure 3a). Furthermore, the differences in temporal variation of the oscillation and stimulus train between amplification and suppression were positively correlated (p = 0.026, Pearson correlation, r = 0.61; Figure 3a), demonstrating that the stimulus evoked change in the stability of the oscillation was mirrored in the pattern of stimulation itself. Such changes in stimulation pattern likely occurred due to the ability of our system to adapt to the length of each individual cycle and/or not stimulate when the phase estimate was deemed unstable (see methods). Further evidence of this brain-machine interaction was seen in the difference in rate (amplifying 25.95 ± 3.34 Hz; suppressing 20.82 ± 2.10 Hz; p = 0.0002, WSRT, W_13_ = 0; Figure 3b) and accuracy (amplifying 73.37 ± 8.48 %; suppressing 45.52 ± 9.09 % within a quarter cycle of target phase; p = 0.0002, WSRT, W_13_ = 0; Figures 3c and S4). These findings suggested that the phase-dependent modulation of spectral power resulted from the equilibrium that developed between the temporal properties of the beta oscillation and the pattern of the stimulus train.

**Figure 3.**
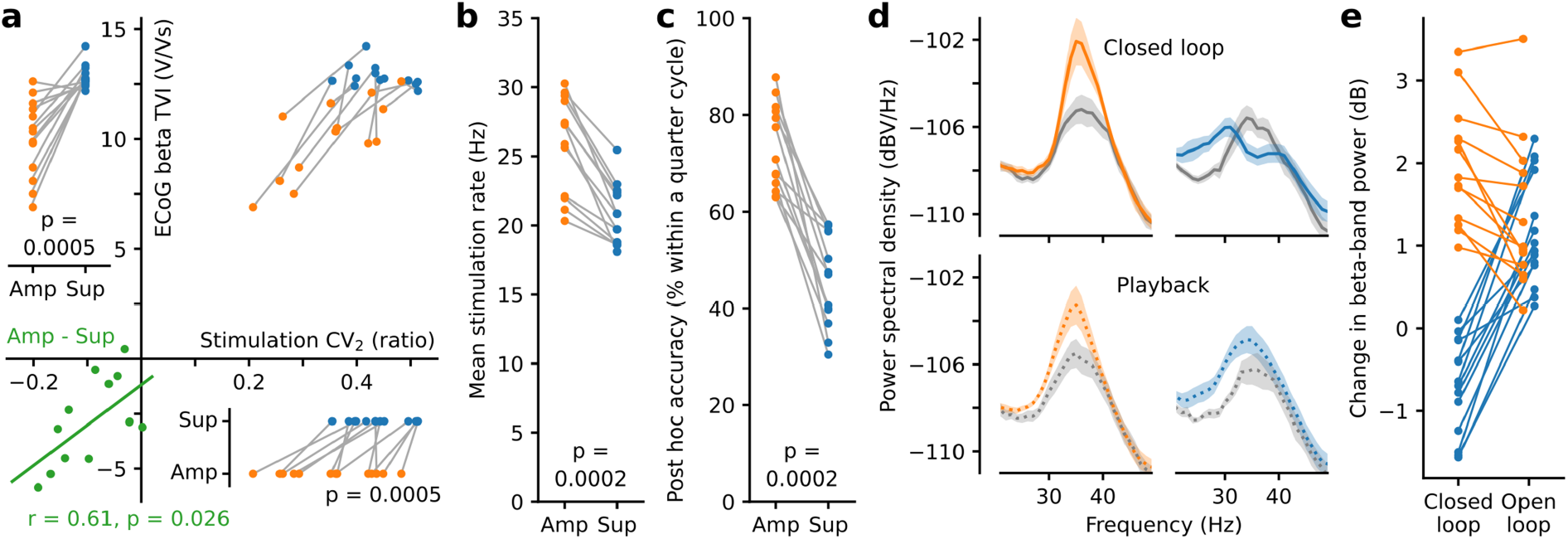
Phase-dependent modulation was produced through brain-machine equilibrium. **a**, Amplified beta-band oscillations were more stable (lower temporal variation; TVI; n = 13 rats) as were their associated stimulation patterns (lower variation in adjacent inter stimulus intervals; CV_2_) reflecting the different state of brain-machine equilibrium associated with amplifying as opposed to suppressing stimulation. The difference between amplifying and suppressing stimulation for both measures was correlated (bottom left). The different states of brain-machine equilibrium also resulted in differences in **b**, mean stimulation rate and **c**, post-hoc accuracy of stimulation phase. **d**, Power spectra from open-loop playback (bottom) of the closed-loop stimulation patterns (n = 27 sessions from 3 rats). **e**, Open loop “playback” of both suppressing and amplifying closed-loop stimulus trains led to beta amplification, but this was lower in magnitude than closed-loop amplification.

### The differences in stimulation pattern alone were not sufficient to reproduce the differences in modulation

To test whether the pattern of stimulation alone could result in phase-related power changes, we delivered closed-loop stimulation at amplifying and suppressing phases then immediately used the recordings to deliver open-loop “playback” of the same stimulation trains (n = 27 recordings, 3 rats). While closed-loop stimulation led to significantly different beta power at amplifying and suppressing phases (p = 1.1e-05, Mann-Whitney rank test, U_13, 14_ = 182), open-loop “playback” did not (p = 0.75, Mann-Whitney rank test, U_13, 14_ = 98; Figures 3d and e). Thus, the temporal relationship between signal and stimulation train was necessary for bidirectional modulation of beta power. Indeed, during open-loop playback both stimulation trains increased beta-band power (previously amplifying p = 0.0002, WSRT to zero, W_13_ = 0; previously suppressing p = 1.2e-04, WSRT to zero, W_14_ = 0; Figure 3e). It is additionally noteworthy that closed-loop amplification was greater than that produced by open-loop playback (p = 0.0007, WSRT, W_13_ = 2; Figure 3e). Overall, the markedly different spectral modulation produced by closed-loop stimulation highlights the importance of the feedback loop.

### Sustained beta amplification slowed movement speed by altering the mode of locomotion

Finally, we sought to establish if the neurophysiological differences seen when targeting stimulation to an amplifying as opposed to a supressing phase were functionally relevant. Sustained beta oscillations are associated with slowing or holding of ongoing movement (Engel and Fries, 2010). We hypothesised that the different modes of beta modulation generated by our system would lead to accompanying changes in behaviour. To test this, speed and gait parameters were evaluated while a subset of GPe and STN stimulated animals (n = 7) traversed a linear track. Amplification, as opposed to suppression, of cortical beta power significantly reduced the speed at which animals moved across the linear track (p = 0.016, WSRT, W_7_ = 0; Figures 4a and S5). The cause of this speed change was a reduction in the ratio of faster bounding-based movement to slower alternating-based movement, irrespective of the animal’s affinity for bounding without stimulation (Figures 4b-e; speed difference correlated with difference in % bounding p = 0.035, Pearson correlation, r = 0.79). Sustained amplification therefore elicited a change in the mode of locomotion, leading to slower movement.

**Figure 4.**
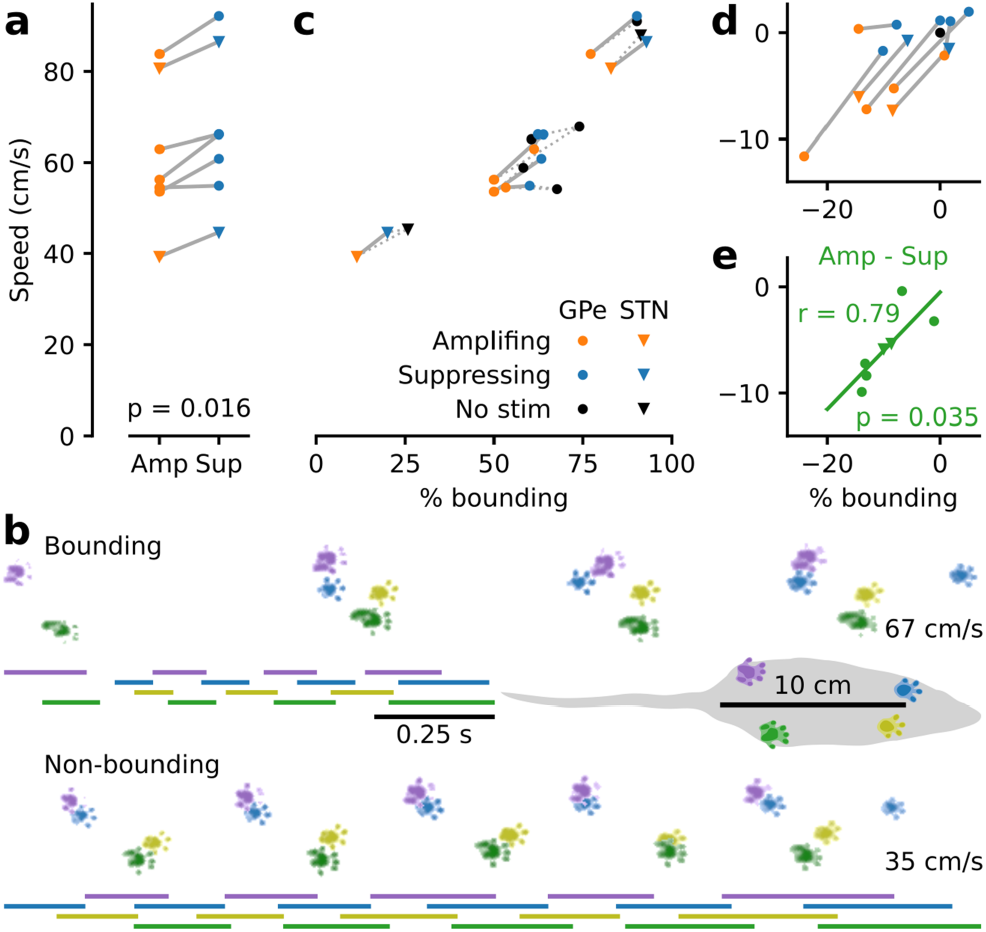
Sustained beta amplification slowed movement speed by altering the mode of locomotion. **a**, Rats (n = 7) traversed a linear track with reduced mean speed in the presence of amplifying stimulation. **b**, Same day, same rat example runs (no simulation; one bounding, one non-bounding) visualised from above. Bars depict the time each foot spend in contact with the track. Average speed is noted on the right. The rat covers a similar distance in half the time when bounding. **c**, The percentage of bounding runs differed across animals and largely determined mean speed. **d**, Same as c plotted relative to the no stimulation condition. **e**, The difference in speed between amplifying and suppressing stimulation correlated with the difference in the percentage of bounding runs.

## Discussion

Cortical circuits entrain widespread populations of neurons across subcortical nuclei, which project directly or indirectly back to the same cortical areas. Such connectivity has been proposed to underlie the generation of beta frequency synchronisation across cortico-basal ganglia circuits (Brazhnik et al., 2016; Cagnan et al., 2019; Mirzaei et al., 2017). Delivering phase-locked stimulation that exploited the temporal relationship between cortical and basal ganglia populations was sufficient to fundamentally change the power and stability of the cortical oscillation. These changes were mirrored by the pattern of the stimulus train, thus an equilibrium emerged, determined by the fixed parameters of the system and the properties of the brain network. Furthermore, altering the phase of stimulus delivery was sufficient to adjust the point of equilibrium. Open-loop playback of the suppressing stimulus train led to *amplification* of the cortical beta oscillation, presumably due to the oscillatory pattern of the stimulus. This reversal in effect clearly illustrates the temporal dependence of signal and stimulus in preventing the oscillation emerging. In contrast, stimulation at the amplifying phase reinforced the stability of the ongoing oscillation, more than could be achieved by the stimulus pattern alone. The system therefore worked like an external node, simultaneously adapting to and influencing the state of the network.

The change in locomotion between amplifying and supressing stimulation – where the only difference between the two conditions was the phase to which the stimulation was targeted – demonstrates that the different states of equilibrium resulted in functionally relevant changes in brain state. Amplification influenced the locomotive preference towards walking over bounding, yet the inter-animal variation in bounding and walking still occupied a similar range to the no stimulation condition. This suggests that the stimulation did not completely disrupt normal function, but instead biased the natural operation of the network, resulting in a shift in behavioural tendencies. Beta oscillations in cortico-basal ganglia networks are associated with a holding of the current motor state (Brittain and Brown, 2014; Engel and Fries, 2010), rather than with a specific type of movement or behaviour. Indeed, the patterned limb movements needed for walking or bounding are mostly coordinated in the brainstem and spinal cord (Ferreira-Pinto et al., 2018). As the rats turned in the reward arena prior to entering the track, they could only move using a walking-like locomotor pattern. Increased beta activity in cortico-basal ganglia networks potentially served to maintain the walking motor program, reducing instances of clean transition into bounding across the track. This ability to bias ongoing behaviour may be particularly advantageous for proposed translational applications of phase-dependent stimulation, where the goal is to shepherd dysfunctional networks away from pathological cognition and behaviour (Takeuchi and Berényi, 2020; Widge and Miller, 2019).

This work supports the general idea that integrating an external system within the intrinsic dynamics of a pathophysiological neural circuit can provide a “network prosthesis”, whereby the closed-loop system compensates for the myriad of maladaptive changes that prevent the network from functioning within its normal range. The approach is clinically tractable by combining the fast real-time algorithm used here with a next generation deep brain stimulation device (Gilron et al., 2021; Opri et al., 2020; Toth et al., 2020). Moreover, the ability to manipulate network oscillations bidirectionally and within their normal functional range enables experiments to define the role of these activities in memory, sleep and other fundamental brain functions.

**Figure S1.**
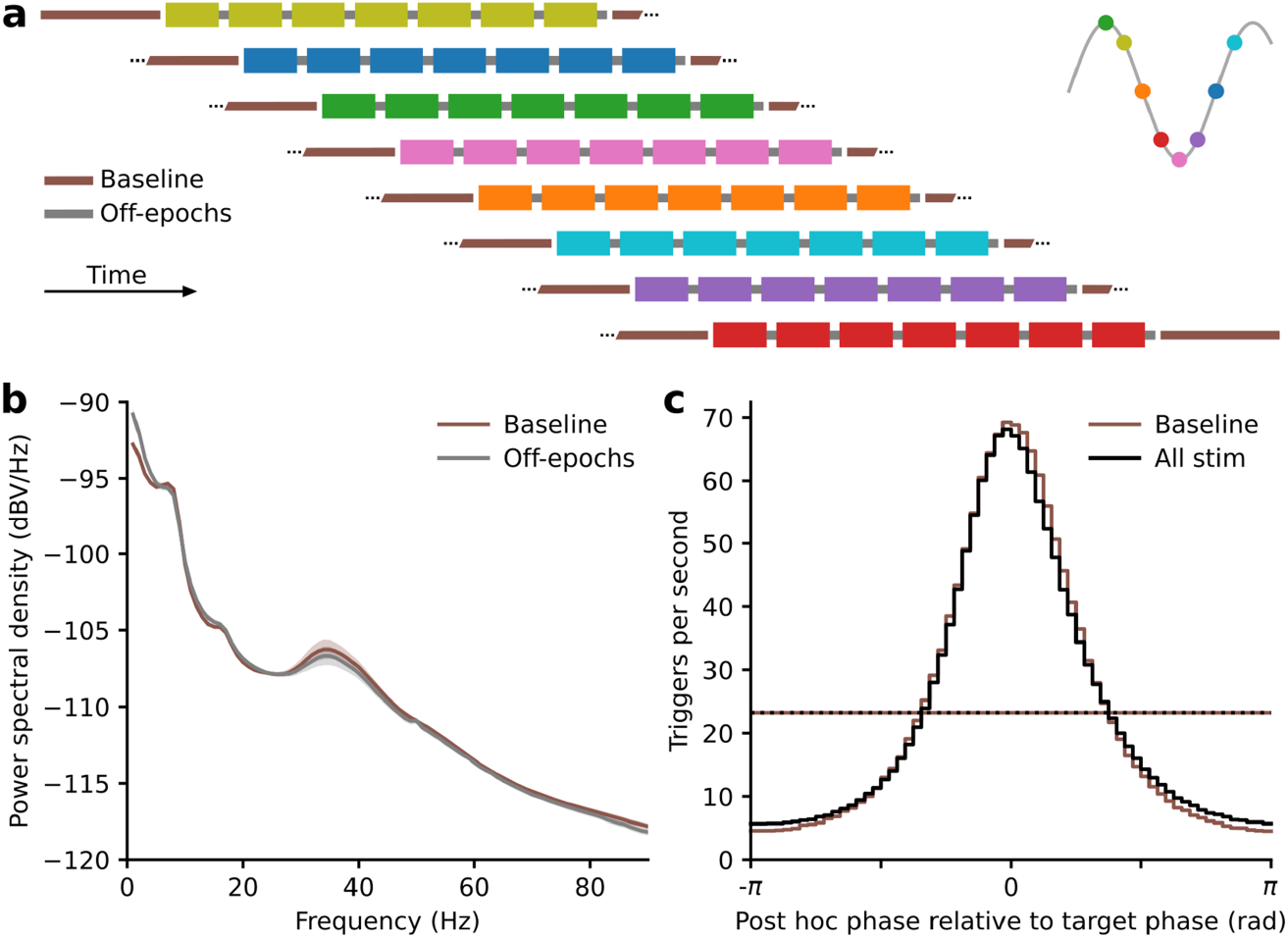
Phase-dependent stimulation targeting parkinsonian beta oscillations. **a**, Schematic of a recording block. Closed loop stimulation occurred during 20 second epochs (on-epochs; colours) separated by 5 second trigger free epochs (off-epochs, grey). Baseline periods (brown) without stimulation were also recorded between each stimulation recording. Target phase (colours, see insert) order was randomised across blocks. **b**, Power spectral density plot (mean ± SEM, n = 13 rats) showing a clear peak in beta-band power in the absence of stimulation recorded from frontal cortical ECoG screws located ipsilateral to the 6-OHDA lesion. Elevated beta-band power in off-epochs (grey) was similar to that seen in baseline recordings (brown). **c**, Trigger accuracy histogram of ECoG phase calculated post hoc plotted relative to target phase. The accuracy and mean trigger rate (dotted lines) of the real-time generated triggers, without stimulation enabled (brown) and with stimulation delivered to the globus pallidus (black) were almost identical, demonstrating that the system was able to maintain full accuracy in the presence of electrical artefacts and physiological perturbation produced by stimulating. Note, post-hoc phase analysis uses future information to calculate phase (primarily due to the band pass filter) and thus by definition represents a different measure to real-time phase regardless of the real-time algorithm used.

**Figure S2.**
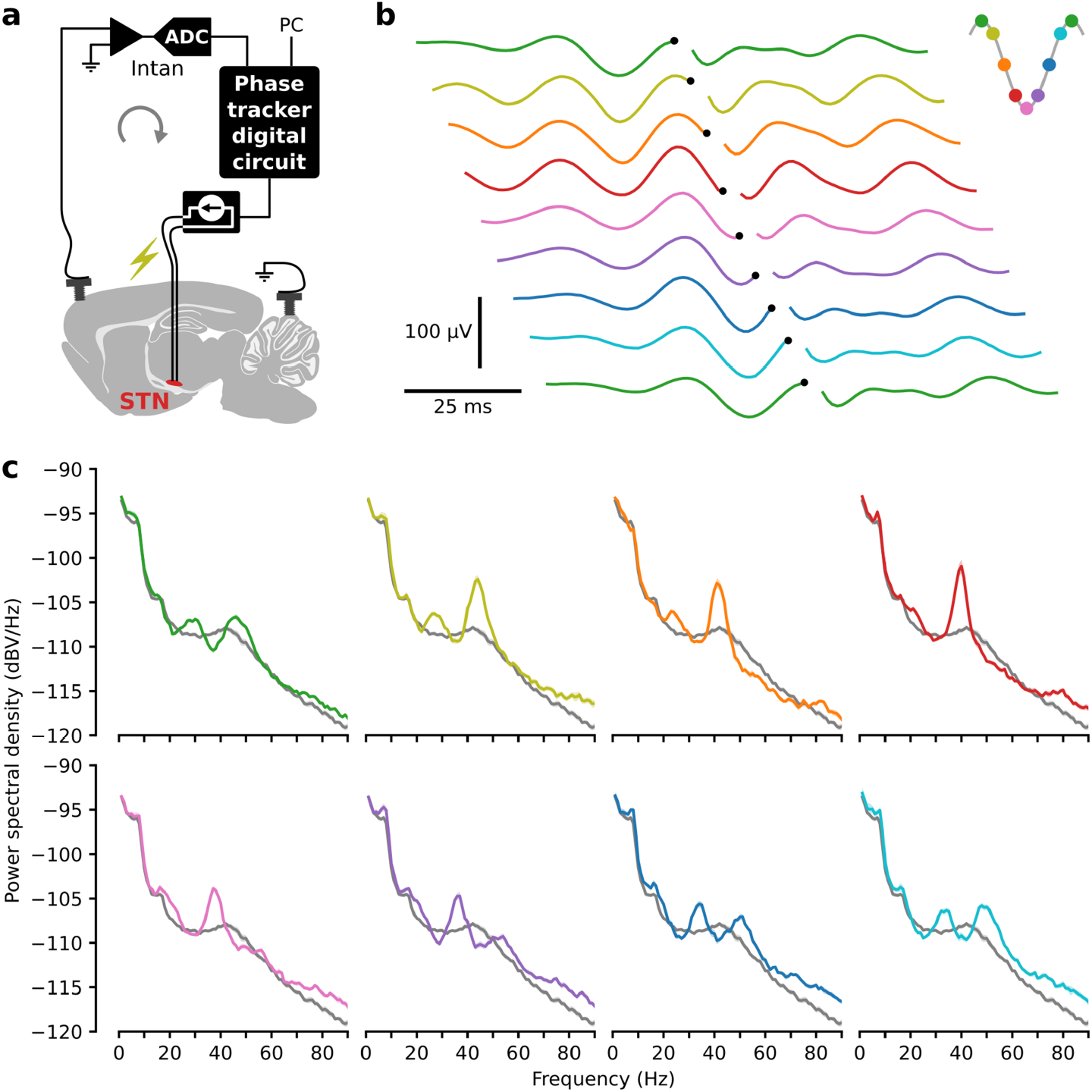
Phase-dependent modulation through stimulation of the subthalamic nucleus. **a**, A digital circuit triggered delivery of electrical pulses to the subthalamic nucleus at a predetermined ECoG beta-band phase. **b**, Stimulation triggered averages from an example block of ECoG recordings. Stimulation was targeted to eight equally spaced phases across separate recordings. To aid visualisation, trigger times (black dots) are staggered across traces and the first trace is repeated. **c**, Power spectra from stimulation on-epochs for each target phase (colours as in **b**) from the same example recordings and for all off-epochs embedded in those same recordings (grey).

**Figure S3.**
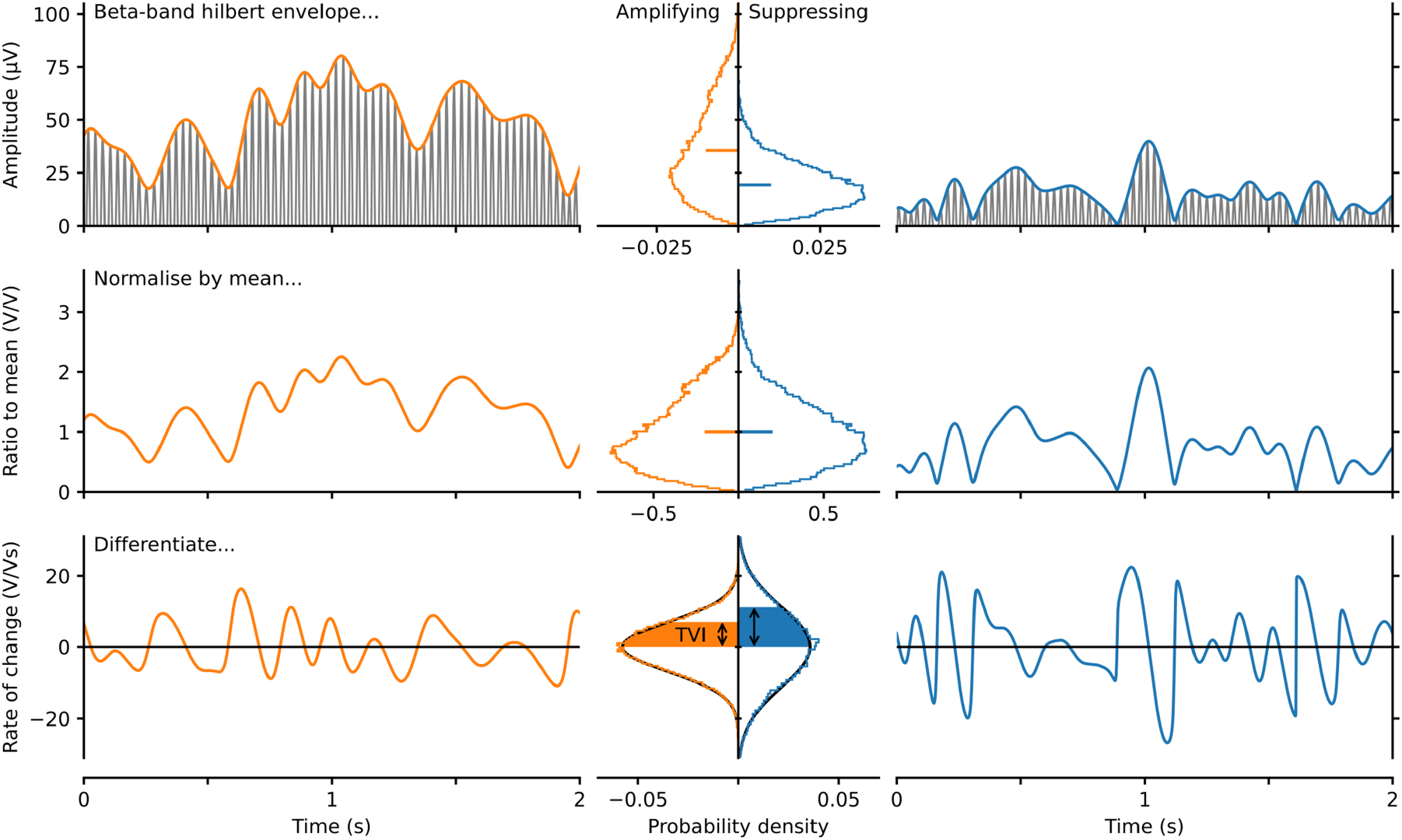
Temporal variation index (TVI). We developed the TVI to provide a measure of beta-band amplitude stability over time independent of differences in mean amplitude. The left (orange) shows the calculation performed on a two-second portion from a recording with stimulation targeted to an amplifying phase whereas the right (blue) is from a suppressing recording. The resulting distributions from the full recording in each case are shown in the middle column. The Hilbert amplitude envelope (top) was calculated from the beta-band filtered signal (grey) resulting in different mean amplitudes. Normalising by the mean resulted in similar amplitude distributions (middle). The standard deviation of these distributions is the coefficient of variation (CV). Here, this provides a normalised measure of variation in amplitude, not variation in time (as is the case when the same measure is applied to time intervals such as neuronal action potential firing). To provide a measure of the fluctuations in amplitude over time we calculated the rate of change (differential) of the amplitude envelope before calculating the standard deviation (bottom). Hence, we defined the TVI as the standard deviation of the derivative of the Hilbert amplitude envelope divided by the envelope mean. It is equivalent and slightly computationally advantageous when calculating the TVI to divide by the mean after differentiation and calculation of the standard deviation. As shown, the derivative results in normally distributed values centred on zero allowing the standard deviation to be fully descriptive of the resulting variation. Here, amplifying stimulation produced a smaller TVI than suppressing stimulation indicating a more stable (less variable) oscillation amplitude.

**Figure S4.**
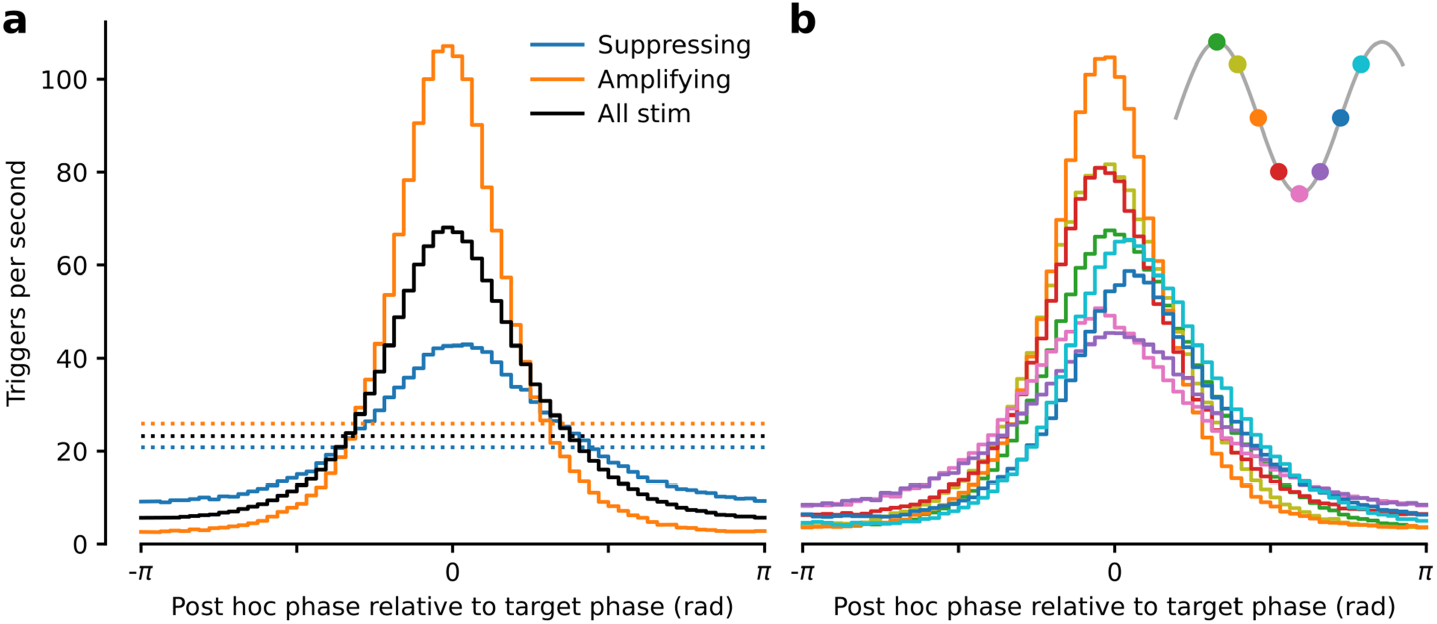
Stimulation rate and accuracy reflected the direction of beta power modulation. **a**, Trigger accuracy histogram of ECoG phase calculated post hoc plotted relative to target phase. Stimulation targeted to the most amplifying phase (orange) produced a higher rate of stimulation on the correct phase (0 rad) and a higher mean stimulation rate (dotted lines) compared to the mean across all target phases (black). Stimulation targeted to the most suppressing phase (blue) led to reduced accuracy and a lower mean stimulation rate (dotted lines) reflecting the systems success in suppressing the oscillation. **b**, Mean trigger accuracy histograms for each target phase plotted relative to target phase with inset showing the target phase of each trace.

**Figure S5.**
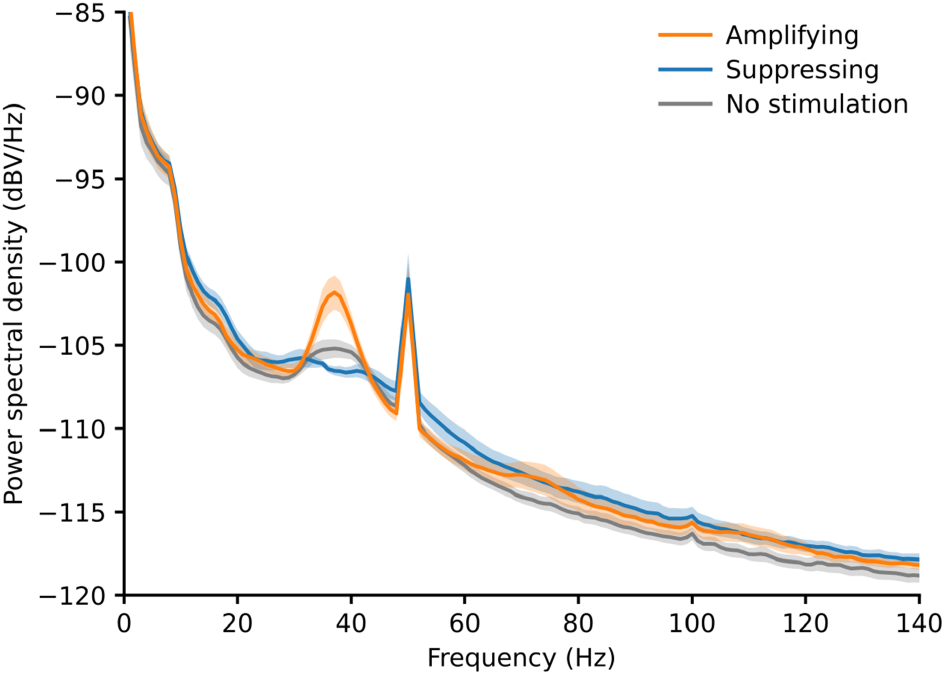
Power spectra from linear track gate analysis experiments. Amplifying, suppressing and no stimulation power spectral density plots (mean ± SEM, n = 7 rats). The spike at 50 Hz is due to interference from the mains power.

## Methods

### Subjects and surgical procedures

Experiments involving animals were conducted in accordance with the UK Animals (Scientific Procedures) Act 1986 under personal and project licenses issued by the Home Office following ethical review. Male Lister Hooded rats (starting weight 350 - 450 g) were housed with free access to food and water in a dedicated housing room with a 12/12-h light/dark cycle and underwent two separate recovery stereotaxic surgical procedures performed under deep anaesthesia using isoflurane (4% induction, 2-0.5% maintenance) and oxygen (2 l/min). Local anaesthetic (Marcaine, 2 mg/kg, 2.5 mg/ml) and non-steroidal anti-inflammatories (Metacam, 1 mg/kg, 5 mg/ml) were administered subcutaneously at the beginning of all surgeries, while opioid analgesia (Vetergesic, 0.3 mg/ml, 0.03 mg/kg) was also provided for three consecutive post-operative days.

The first surgical procedure produced a unilateral lesion of dopaminergic neurons of the substantia nigra pars compacta by intracranial injection of the neurotoxin 6-hydroxydopamine (6-OHDA) at their cell bodies through a glass pipette located 4.9 mm posterior and 2.3 mm lateral from bregma at a depth of 7.8 mm from the brain surface. 6-OHDA was dissolved immediately before use in phosphate buffer solution containing 0.02% w/v ascorbate to a final concentration of 6 mg/ml. Between 0.10 and 0.15μl of 6-OHDA solution was injected at a rate of 0.01μl/min through the pipette, which was left in place a further 5 minutes before being withdrawn. Front-foot use asymmetry pre- and post-lesion while rearing against the wall of transparent acrylic cylinder was used to indicate lesion severity and select candidates for electrode implant.

Following full recovery and not less than 13 days later, selected animals underwent a second surgical procedure to implant two pairs of stainless steel stimulation electrodes (California Fine Wire, stainless steel, bifilar, heavy formvar insulation, 127μm strand diameter). Electrodes were secured with bone cement and six M1.4 × 3 mm stainless steel screws. For stimulation of the globus pallidus (GPe), electrodes were implanted 1 mm posterior and 3.1 mm lateral from bregma at a depth of 6 mm from the brain surface. For stimulation of the subthalamic nucleus (STN) electrodes were implanted 3.8 mm posterior and 2.5 mm lateral from bregma at a depth of 7.2 mm from the brain surface. ECoG was measured from the most frontal screw located above motor cortex at approximately 4.6 mm anterior and 1.6 mm lateral from bregma referenced to two screws above cerebellum. Neurotoxin injections, electrical stimulation and ECoG recording were all performed on the right hemisphere. The position of stimulation electrodes was examined post-mortem. All animals included in the study had stimulation electrodes in or adjacent to the GPe and/or STN.

Closed-loop stimulation, data acquisition and behavioural testing took place in a separate room during dedicated recording sessions spaced over multiple weeks. Upon completion, rats were deeply anesthetized with isoflurane (4%) and pentobarbital (3 ml, Pentoject, 200 mg/ml) and transcardially perfused with phosphate-buffered saline (PBS) followed by fixative (paraformaldehyde dissolved in PBS, 4%, wt/vol). Brains were extracted and sectioned. Lesions were verified using antibodies to tyrosine hydroxylase (rabbit anti-TH primary; diluted 1:1000; Millipore Cat# AB152, RRID:AB_390204; donkey anti-rabbit Alexa Fluor 488 secondary; diluted 1:1000; Thermo Fisher Scientific Cat# A-21206, RRID:AB_2535792) to visualise the unilateral loss of cell bodies in the SNc and loss of dopaminergic innervation in the dorsal striatum.

### Open field eight phase protocol

Rats freely explored a 90 by 50 cm dimly lit open field surrounded on three sides, above and below with electrical shielding with a black curtain for access on the remaining side. Three to six recording blocks were performed per rat. Recording blocks consisted of eight pairs of recordings with a different real-time target phase applied in each pair (Figure S1a). Target phase order was randomised across blocks. Stimulation was enabled in the first recording of each pair (stimulation recordings) and disabled in the second (baseline recordings). Stimulation recordings were approximately 4.5 minutes in duration and baseline recordings were approximately 2 minutes in duration. The real-time system generated triggers during 20 second epochs (on epochs) separated by 5 second trigger free epochs (off epochs) and was in operation across all recordings. Recordings were made both when triggers were used to drive stimulation and with stimulation disabled. Generating real-time triggers in the absence of stimulation during baseline recordings allowed comparison of the real-time system performance of phase tracking with and without stimulation influencing the neural activity. These recordings also provided the opportunity to assess baseline physiology in the absence of stimulation. Embedding short epochs lacking stimulation within recordings with stimulation provided an internal comparison point to assess the effect of stimulation without the need to correct for changes occurring over longer time frames such as arousal, brain state, general movement levels and external electrical noise. The centre frequency (f_c_) for real-time phase tracking for each block was chosen at the start of the block based on power spectra generated from previous recordings from the same rat and was in the range 35 to 41 Hz.

### Electrocorticogram and electrical stimulation

Electrocorticogram (ECoG) signals measured from M1.4 stainless steel screws referenced to two cerebellar screws were amplified and digitised using a RHD2000 family amplifying and digitising headstage connected to a RHD USB Interface Board both from Intan Technologies (intantech.com). Electrical stimulation was driven using a fully isolated current source (battery powered with an optically coupled trigger, A365RC, World Precision Instruments) with the source and sink terminals connected to the adjacent strands of the pair of stainless steel wires forming the stimulation electrode. Stimulation events were biphasic consisting of two consecutive pulses of opposite polarity each 50, 60 or 70μA in amplitude and 95μs in duration separated by 10μs. To achieve closed-loop stimulation, a custom designed digital circuit was used to access the digital data stream from the headstage, track phase within a band of interest in real-time and generate digital pulses to activate the optically coupled trigger of the stimulation current source. The custom designed digital circuit was implemented using extra available digital circuitry within the field programmable gate array (FPGA) located on the RHD USB Interface Board.

### Real-time phase tracking and closed-loop trigger algorithm

ECoG recordings were performed with a sample rate of 20 kHz per channel to capture the full detail of stimulation artefacts aiding their removal in post hoc analysis. However, for phase tracking in real-time, the data was downsampled by producing a single sample from the sum of 8 to 12 (*N*_*dn*_) successive samples, since tracking beta-band phase did not require such a high number of samples per cycle. Thus, the phase tracking component operated at sample rates in the range 1.67 to 2.5 kHz (varying the sample rate was one of the ways to achieve different target oscillation centre frequencies).

To track phase in real-time, an iterative algorithm was used to perform a continuous transform on the data stream. This allowed a phase estimate to be instantaneously produced corresponding to each value of the downsampled data stream. A bandlimited complex estimate of the signal was generated from the downsampled wideband data stream by, on receipt of each sample, updating a complex weight register that maintained the relationship between the estimate of the signal and a pair of continuously oscillating reference sine and cosine waves. The error between the real part of the estimate signal and the current ADC sample was used to update the weight register for use with the next ADC sample. The centre frequency of the band of interest was set by the frequency of the reference waves. This depended on the downsampled sample rate and the number of values in the lookup table containing a quarter of a sine wave used to generate the reference waves (*N*_*points*_, 11 to 15 values per quarter cycle were used). Thus the centre frequency of the phase tracker pass band was given by (*f*_*c*_ = 20000/*N*_*dn*_/*N*_*points*_/4). The width of the band of interest was set by a constant coefficient on the error term used to update the weight register. This essentially smoothed the reference wave by dampening the rate of convergence between the estimate and the signal. Finally, a continuous phase estimate was produced by calculating the polar angle of the complex signal estimate using a CORDIC (Volder, 1959).

A real-time trigger was generated when the phase estimate passed into the target phase range unless it had been less than 0.8 of a target frequency (*f*_*c*_) period since the most recent passing into the target phase range. This was important to reduce stimulation at times of poor phase estimates caused by low power in the target frequency range. Additionally, a phase shift corresponding to half the stimulation width (100μs) was subtracted from the target phase causing the stimulation to be triggered slightly early allowing the middle of the stimulation to occur at the desired target phase. To reduce the effect of electrical stimulation artefacts on the phase estimate, a digital hold circuit operating on the 20 kHz data stream held the sample preceding the stimulation trigger on the input of the downsampling circuit for 600μs from the start of the trigger. An offset removal digital filter was also implemented on the input to the phase tracking algorithm to remove residual offset remaining from the analogue front-end electronics.

### Data analysis

Post hoc analyses of ECoG spectral properties, stimulation phase and behavioural data including statistical tests were performed using SciPy (scipy.org). Text values are reported as mean ± standard deviation and error bars in plots show mean ± standard error of the mean unless otherwise stated. First stimulation artefacts were removed from the 20 kHz recordings by interpolation between the sample immediately preceding the stimulation and the sample 1.7 ms later. The resulting signals were downsampled to 1 kHz in two steps using finite impulse response anti-aliasing filters (designed using scipy.signal.remez(), combined pass band ripple less than 0.001 dB below 420 Hz, stop band attenuation greater than 90 dB above 498 Hz). All spectra and phase calculations were performed on the artefact-removed and downsampled signal. Post hoc trigger phase in the beta-band was calculated by filtering (scipy.signal.firwin(), numtaps = 513) the signal between ±5 Hz of the centre frequency (f_c_) chosen for real-time phase tracking. The trigger phase was then calculated as the polar angle of the Hilbert transform analytic signal at the midpoint of the biphasic trigger. Power spectral density (PSD) calculations were performed using Welch’s method (Welch, 1967) with a resolution of 1 Hz spectral bins. Changes in beta-band power were calculated from the respective PSDs (mean of f_c_ bin ± 5 bins inclusive, i.e. mean of 11 bins total) in dB. The Watson-Williams test was performed using pycircstat (pypi.org/project/pycircstat). For capturing the variation in stimulation dynamics, coefficient of variation 2 (CV_2_) was used (Holt et al., 1996). It was calculated as the mean value across the stimulation train of two times the absolute value of the difference over the sum of adjacent stimulation intervals. It has a value of one for a Poisson process and zero for regularly spaced (fixed frequency) stimulation.

### Temporal variation index

In order to quantify the interplay between the stimulus train and the beta oscillation, we required a measure of the oscillatory variability in amplitude with respect to time. Coefficient of variation-type measures when applied to time intervals, capture the variability in time for point processes such as neuronal spiking or stimulation trains. However, for constant sample rate processes such as oscillatory amplitude they cannot be directly applied to the time axis and give a measure of the variability in amplitude, not time. Consider two identical signals with one difference; one is progressing at twice the rate of the other. Their variance and coefficient of variation (CV) in amplitude would be identical. However, the variance in their rate of change would differ and this is the property we sought to measure. The more slowly progressing signal could be described as more stable as its amplitude changes more slowly. We thus developed the temporal variation index (TVI; Figure S3) where the differential of the signal is taken before variance is measured. Differentiation of such a signal results in values that are normally distributed being centred on zero and hence can be fully described by their standard deviation (see Figure S3 bottom row). A second requirement was that the measure should be independent of mean amplitude as we wished to compare the temporal dynamics between the amplified and suppressed signals. This was achieved by normalisation by the mean. Without normalisation, signals with a higher mean would have a higher TVI. This normalisation is equivalent to that to produce CV. Thus, we defined TVI as the standard deviation of the derivative of the Hilbert amplitude envelope divided by the envelope mean. A more stable signal will have a lower TVI and a more variable signal will have a higher TVI.

### Linear track gait analysis

Gait analysis during closed-loop stimulation was performed using a customised version of the CatWalk XT (Noldus, Netherlands). The CatWalk system consists of a glass walkway raised above a camera capturing at 100 fps. The glass plate is illuminated by green light that reflects within the glass at points being touched, allowing for semi-automated detection of footprints as rats traverse the walkway. The CatWalk was modified to include arenas extending from the ends of the track each containing a reward port. Aluminium sheeting was attached along the walls of the track to provide electrical shielding and increase the wall height. The red strip light above the track was replaced with red ambient lighting allowing the addition of a zip line to accommodate the recording and stimulation tether.

Sugar pellets were delivered by the experimenter through dedicated tubing to the reward delivery ports at alternating ends of the track. During training, tethered rats were allowed to freely explore the glass walkway and reward areas for approximately four fifteen-minute sessions until they consistently crossed the walkway to collect reward. Amplifying and suppressing stimulation phases and stimulation target (GPe or STN) for each rat were selected based on spectra generated from open field eight phase recordings. During data acquisition, no stimulation, amplifying-phase stimulation and suppressing-phase stimulation were applied in separate recordings in a randomised order. Recordings were between five and twenty minutes in duration and typically consisted of eight valid runs with multiple groups of recordings per animal collected across different days. Gait analysis was performed on rats in which successful amplification and suppression was achieved during linear track recordings. Damaged implants prevented collection of linear track data from two of the 13 rats comprising the open field data. Post-hoc analysis of linear track ECoG data revealed that amplification and suppression of beta-band power did not occur in a further four rendering the data unsuitable for the comparison of amplifying and suppressing stimulation (two had no modulation and the incorrect target phases were selected during data collection for another two). Of the remaining seven included in gait analysis, five received stimulation of the GPe and two received stimulation of the STN.

Paw prints were detected and descriptive properties generated using CatWalk XT software version 10.6.608 (Noldus, Netherlands) and the resulting output further analysed using SciPy (scipy.org). Only runs less than four seconds in duration having at least eight valid footprints and an average speed of greater than 10 cm/s were included in the analysis. To identify faulty recordings, the total spectral power below 200 Hz was calculated for each recording and those in the upper tail of the resulting distribution were excluded (20 of 420). The average number of valid runs per condition per animal was 100.7 with a minimum of 69. Average speed for each run was calculated as the average distance travelled divided by time taken for each step cycle i.e. from start of contact to start of next contact for each cycle of each foot (CatWalk XT). Runs were classified as bounding if each forelimb contact was followed or preceded by the other across the entire valid run with the same being true of hindlimbs e.g. front-front-hind-hind-front-front etc., never front-hind-front (CatWalk XT 100 % cruciate or rotate). The percentage bounding was the percent of total valid runs for that condition that were classified as bounding.

